# Virtual screening and zebrafish phenotype-based evaluation argues against repurposing 4-phenylbutyrate for STXBP1-related disorders

**DOI:** 10.64898/2026.05.07.723632

**Authors:** Aline Frick, Paige Whyte-Fagundes, Scott C. Baraban

## Abstract

Syntaxin-binding protein 1 (*STXBP1*) mutations lead to severe epilepsy, intellectual disability, developmental delay, and movement disorder. Effective treatments for these conditions do not exist. Recent studies in Munc18-1 (*STXBP1*) *C. elegans* models demonstrate that 4-phenylbutyrate (4-PBA) or related pharmacological chaperones stabilize Munc18-1 protein levels and rescue locomotion deficits. These studies suggest a novel treatment strategy for these patients. Here, we used a *stxbp1a* zebrafish model with a profound movement disorder to screen 4-PBA and alternative structural analogs identified using artificial intelligence (AI)-based screening. Automated locomotion assays conducted on larval *stxbp1a* mutant zebrafish at 5 days post-fertilization (dpf) confirm and extend the movement disorder endophenotype. Drug treatment (4-PBA or 16 identified candidates) failed to rescue the *stxbp1a* mutant zebrafish locomotion deficit. Electrophysiology studies in a *stxbp1b* zebrafish model characterized by spontaneous seizure activity (i.e., epilepsy) failed to detect a reduction in ictal-like events with 4-PBA treatment. Taken together, our results suggest caution in repurposing 4-PBA or related compounds for treatment of STXBP1 disorders.

## Introduction

Syntaxin-binding protein 1 (STXBP1) is a core component of a highly conserved protein complex governing exocytosis e.g., soluble N-ethylmaleimide-sensitive factor attachment protein receptors (SNARE) complex (Han et al., 2017). This complex controls machinery necessary for presynaptic neurotransmitter release and plays a critical role in neural development, synaptic plasticity and cognitive function (Jahn and Fasshauer, 2012, Zhou et al., 2017, Chen et al., 2021). Heterozygous *de novo* mutations in *STXBP1* are associated with severe childhood epilepsy, intellectual disability, autism and movement disorders (Stamberger et al., 2016, Abramov et al., 2021). Initially identified as a missense mutation causing Ohtahara syndrome (Saitsu et al., 2008), a developmental epileptic encephalopathy, missense, nonsense, truncation, frame shift or microdeletion *STXBP1* mutations are also linked to West (Otsuka et al., 2010), Lennox-Gastuat (Epi et al., 2013), Dravet (Carvill et al., 2014) and Rett (Romaniello et al., 2015) syndromes. Seizures, usually within the first year, are a common feature of these neurological conditions. *STXBP1* mutations in patients with ataxia or movement disorders have also been identified (Gburek-Augustat et al., 2016). A recent study of 162 *STXBP1*-related disorder patients revealed delay in gross motor movement milestones as a primary phenotype in nearly half of these individuals (Xian et al., 2023). In all these conditions available treatment options (e.g., antiseizure medications such as vigabatrin, levetiracetam, phenobarbital or clobazam) are limiting and ineffective. Morbidity and mortality incidence rates three-fold higher than the general epilepsy population (Furia et al., 2024) are also common. A lack of adequate therapies to address complex seizures and comorbidities seen in this population spurred recent interest in antisense oligonucleotide (ASO), CRISPR activation, and antisense long non-coding RNAs (SINEUPs). Despite potentially transformative potential for these types of therapies for rare disease communities, over 90% of marketed therapies are small molecule drugs (Gurevich and Gurevich, 2014).

4-phenylbutyrate (4-PBA) was identified by Burre and colleagues as a chemical chaperone which could stabilize protein expression and prevent misfolded protein aggregation in a *C. elegans* Munc-18 (Stxbp1) model characterized by lack of movement (Guiberson et al., 2024, Abramov et al., 2021, Guiberson et al., 2018). Similarly, Kang and colleagues describe absence-like spike-wave electroencephalographic (EEG) activity in *Slc6a1*^*+/-*^ and *Gabrg2*^*+/Q390X*^ mice that was reduced following 4-PBA treatment (Shen et al., 2024, Nwosu et al., 2022). Approved for treatment of urea cycle disorders, 4-PBA was also studied for neurological conditions such as spinal muscular atrophy (Mercuri et al., 2007) and amyotrophic lateral sclerosis (Ketabforoush et al., 2024) but efficacy was not established with either trial. Based on these observations, 4-PBA as a potentially repurposed drug candidate (e.g., an approved drug used for a different medical condition for which it was intended) may offer a streamlined path to clinical application for STXBP1-related disorders. Indeed, open-label studies have begun to evaluate safety and tolerability of 4-PBA in a small cohort of STXBP1 patients (Grinspan ZM, 2024). While results of these compassionate-use studies are mixed, a drug repurposing approach represents an efficient strategy to reduce both time and cost of new therapy development. A strategy that successfully contributed to Food and Drug Administration (FDA) approved treatment options for Tuberous Sclerosis Complex (everolimus) and Dravet syndrome (fenfluramine)(Schoonjans and Ceulemans, 2019, Johannessen Landmark et al., 2021).

The emergence of haploinsufficient preclinical mouse, worm and zebrafish models for STXBP1-related disorders further accelerated research for new treatments. Among these, zebrafish (*Danio rerio*) offer a powerful platform for efficient *in vivo* drug screening (Shcheglovitov and Peterson, 2021, Patton et al., 2021, Parng et al., 2002). In contrast to *C. elegans* or *Drosophila melanogaster*, they are small vertebrates with similar tissues, development and physiology to humans. In phenotype-based drug screens, candidates can be identified for their potential to ameliorate a disease phenotype independent of known or predicted targets. Advantages of zebrafish for *in vivo* phenotype-based screening also include high conservation of genes (Howe et al., 2013) and protein similarity with human, rapid development, and large clutch sizes offering statistical power for high-throughput screening (HTS) efforts. To date more than seventy HTS small-molecule screens in zebrafish have been reported, with successful identification of approximately a dozen drug candidates currently in clinical trials (Dash and Patnaik, 2023, Rosa et al., 2022, Patton et al., 2021). Among them, a first-generation antihistamine clemizole (Finkelstein et al., 1960), which is currently in a randomized, double-blind, placebo-controlled clinical trial for Dravet syndrome (ARGUS, ClinicalTrials.gov ID: NCT04462770), was identified in a phenotype-based screen using a *scn1a* zebrafish model (Baraban et al., 2013). This model also successfully predicted (Whyte-Fagundes et al., 2024) the Phase 3 SKYLINE clinical trial failure of soticlestat (Sullivan et al., 2026). Although drug repurposing traditionally relies on experimental approaches testing thousands of compounds from commercially available libraries, rapidly evolving computational approaches offer a strategy for virtual screening of libraries to identify a subset of drugs for phenotype-based assays.

Here, we employed artificial intelligence (AI) technology to accelerate drug repurposing for STXBP1-related disorders. Our approach was based on a parameter-free online server program, [http://cao.labshare.cn/drugrep/, (Gan et al., 2023)] that offers a ligand-based virtual screening platform measuring the similarity between a submitted ligand and compounds in an approved drug library. Following on preclinical and early compassionate-use interest, we used 4-PBA as a ligand and performed AI-based virtual screening with a Computer-Aided Drug Design (CADD) program available in DrugRep. Virtual screening identified candidates (Table 1). Then we conducted *in vivo* phenotype-based locomotion assays with a *stxbp1a* mutant zebrafish line characterized by a profound lack of movement. In parallel studies to evaluate potential therapeutic efficacy against seizures, we tested 4-PBA in epileptic *stxbp1b* mutants using local field potential (LFP) recordings. STXBP1-related phenotypes were not improved with 4-PBA or any related drug candidates in these zebrafish assays.

**Table 1:**
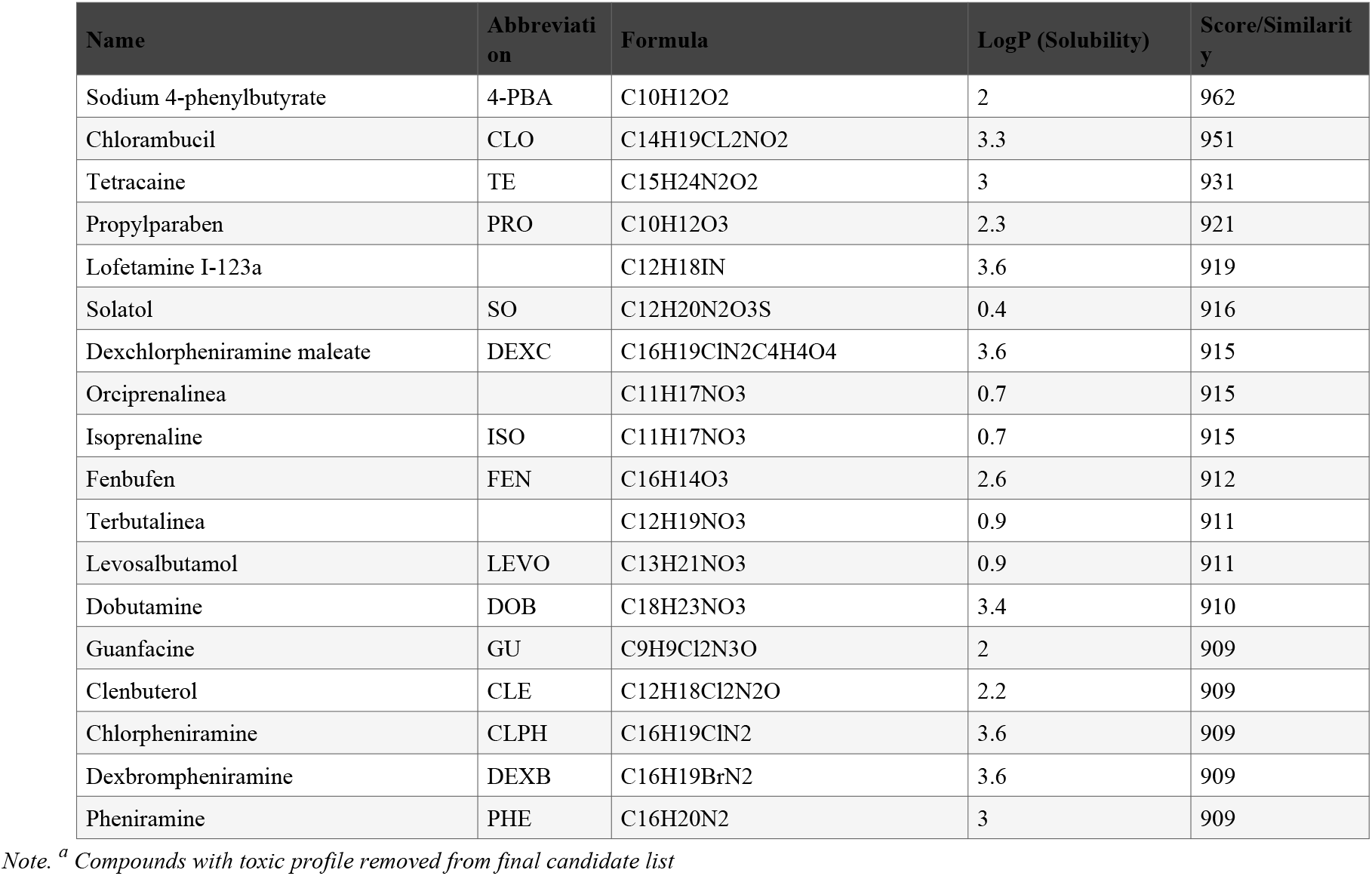
List of virtual candidates using Flexi-LS method.

## Results

Preclinical animal models are vital tools in identification and development of novel therapies. Here, we used CRISPR/Cas9-generated zebrafish (i) carrying a 4 base-pair frameshift deletion in *stxbpb1a*, a paralog sharing 87% amino acid sequence identity with human or (ii) carrying a 12 base-pair loss-of-function deletion in *stxbpb1b*, a paralog sharing 79% amino acid sequence identity with human (Grone et al., 2016). At 3 dpf homozygous *stxbp1a* mutant zebrafish larvae fail to hatch out of the chorion; homozygous *stxbp1b* mutant zebrafish larvae exhibit normal development.

To evaluate potential therapeutic efficacy, we focused on a robust behavioral comorbidity observed in *stxbp1a* zebrafish e.g., lack of spontaneous movement. Using an automated locomotion tracking system (DanioVision, Noldus Technology Inc.) that uses single-point detection to track objects darker than background we first obtained locomotion data from wild-type (WT) and *stxbp1a* larvae at 5 dpf. As expected (Grone et al., 2017), *stxbp1a* zebrafish swim movement was significantly reduced by 73% compared to WT control, as measured by total distance moved. Next we used tetracaine (200 M) as a negative control to establish endogenous ‘noise’ levels for automated single-point detection locomotion tracking. Tetracaine reduced WT movement by 69% compared to control (**Fig. 1A; Supplemental Table 1**), consistent with anesthesia. Notably, ‘total distance traveled’ values were similar for *stxbp1a* compared to tetracaine-treated WT larvae indicative of a severe movement deficit. Next, to infer an upper locomotion threshold we tested whether drugs reported to induce hyper-locomotor activity in zebrafish - MK-801 (Chen et al., 2010, Seibt et al., 2010) and caffeine (Lovin et al., 2023) alter total distance traveled in *stxbp1a* or WT zebrafish at 5 dpf. At bath concentrations that did not cause toxicity, neither MK-801 (25 or 50 M) nor caffeine (25 or 50 M) increased locomotion for *stxbp1a* or WT larvae (**Fig. 1B**). However, a decrease in total distance traveled was observed at 25 M MK-801 in WT larvae consistent with the decrease in average distance travelled at this concentration reported by Li and colleagues (Li et al., 2018).

**Figure 1.**
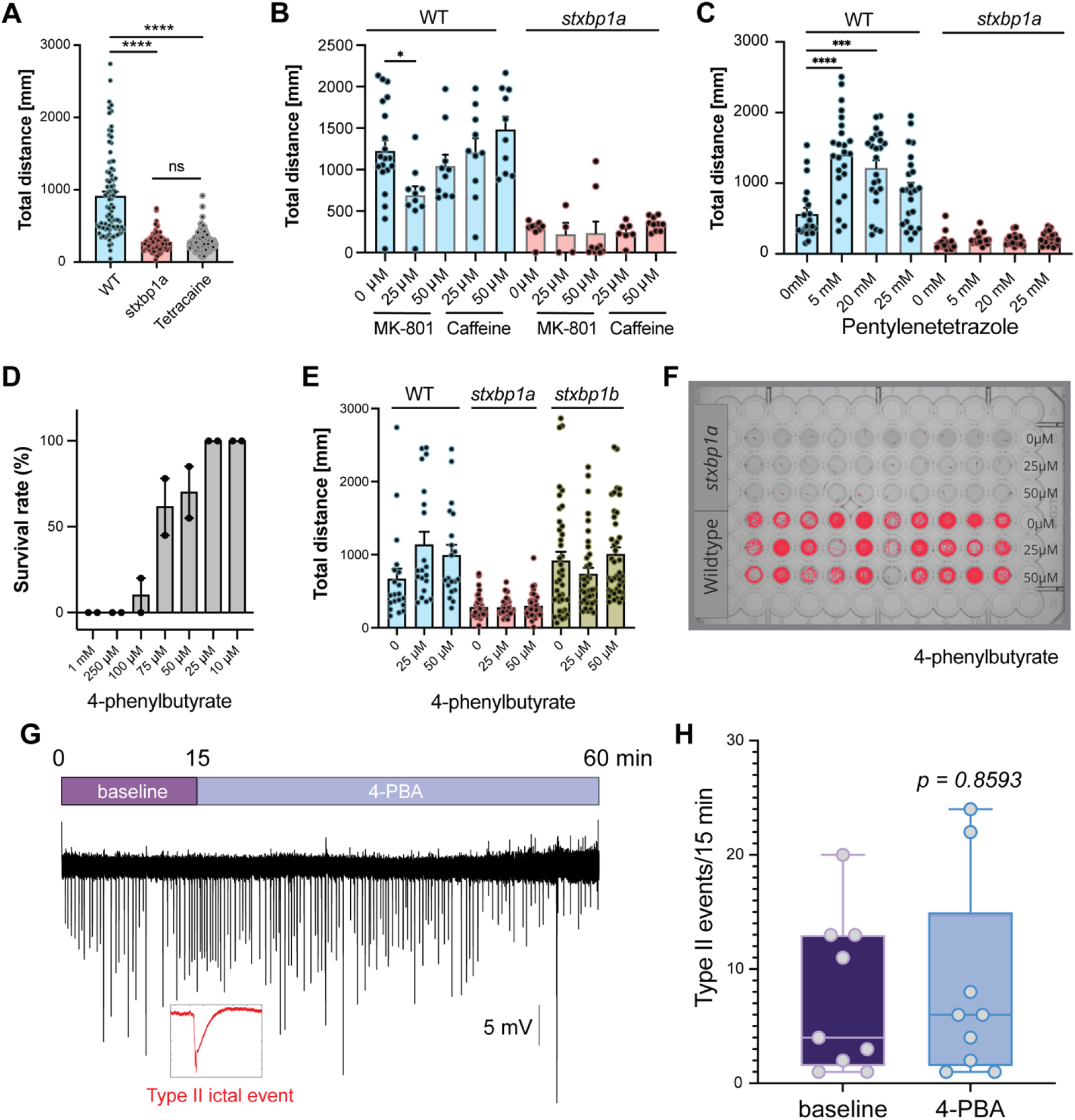
Movement deficit and response to 4-PBA in *stxbp1* mutant larvae. (**A**) Bar plots of total distance traveled during a 10 min tracking epoch. Data are shown for age-matched WT, *stxbp1a*^*-/-*^ in normal embryo media (*stxbp1a*) and *stxbp1a*^*-/-*^ in normal embryo media with tetracaine (tetracaine) larvae. Data is shown as mean ± S.E.M. ****significance p < 0.0001One-way ANOVA. (**B**) Bar plots of total distance traveled during a 10 min tracking epoch. Data are shown for *stxbp1a*^*-/-*^ and WT larvae exposed to MK-801 or caffeine. Data is shown as mean ± S.E.M. (**C**) Bar plots of total distance traveled during a 10 min tracking epoch. Data are shown for *stxbp1a*^*-/-*^ and WT larvae exposed to pentylenetetrazole. Data is shown as mean ± S.E.M. Significant differences are denoted as ****p≤0.0001, ***p≤0.001, **p≤0.01. One-way ANOVA with Dunnett’s multiple comparison test. (**D**) Bar plot of the percent of live, healthy WT larvae following exposure to 4-phenylbutyrate for 48 hr. Note that 100% of larvae survive exposures at 10 and 25 µM. N = 20-40 fish for each condition. (**E**) Bar plots of total distance traveled during a 10 min tracking epoch. Data are shown for age-matched WT, *stxbp1b*^*-/-*^ and *stxbp1a*^*-/-*^ larvae exposed to normal embryo media (0) or 4-phenylbutyrate at 10 and 25 µM. N=20-30 fish in each concentration with 2-3 replicates (**F**) Representation of cumulative tracking of the spontaneous locomotion of 5 dpf WT and *stxbp1a* mutant larvae for 20 minutes after 48 hours of incubation in 4-PBA. Each well represents an individual fish. Movement is shown in red. (**G**) Representative 60◼min local field potential (LFP) trace for *stxbp1b* mutant zebrafish at baseline (0-15 min) and continuous bath exposure to 50 M 4-PBA (45–60). Inset shows a high-resolution image of a typical Type II ictal-like seizure event. Significance *p* = 0.8593 Unpaired t test, 95% confidence interval = -7.18 to 8.51 (**H**) Individual value plot of ictal event frequency measured at baseline (0–15 min) and 45–60 min after continuous bath application of 4-PBA.

Because larval seizure activities are consistently associated with increased swim movement (Baraban et al., 2005, Shaw et al., 2022, Afrikanova et al., 2013), we next tested a chemoconvulsant (pentylenetetrazole, PTZ) in *stxbp1a* and WT larvae at 5 dpf. PTZ also failed to elicit an increase in total distance traveled at the same concentration range (5 – 25 mM) in *stxbp1a* mutants whereas movement was significantly increased in WT zebrafish (**Fig. 1C**). Taken together, these studies confirm and extend observations of a movement deficit comorbidity in *stxbp1a* mutant zebrafish.

Humans, mice, worms, and zebrafish with *STXBP1* mutation exhibit locomotion/movement deficit. In homozygote Munc18-1 mutant worms (*C. elegans*), 5 mM 4-PBA rescued this locomotion deficit (Guiberson et al., 2024). To adapt this protocol to larval zebrafish we first performed studies to determine Maximum Tolerated Concentration (MTC); 4-PBA was toxic to larval zebrafish at concentrations above 50 M (**Fig. 1D**). We then used locomotion assays to test two 4-PBA concentrations in WT, *stxbp1b*^*-/-*^ and *stxbp1a*^*-/-*^ zebrafish. 4-PBA exerted no effect on normal WT and *stxbp1b* movement and failed to rescue the movement deficit seen in *stxbp1a* mutant larvae at 25 or 50 M (**Fig. 1E**). Representative locomotion tracks are shown for *stxbp1a*^-/-^ and age-matched WT larvae in **Fig. 1F**. Because *stxbp1b* mutants exhibit an epilepsy phenotype characterized by spontaneous ictal-like activity in local field potential (LFP) recordings (**Fig. 1G**) we next tested whether 4-PBA could rescue this endophenotype. Recording ictal-like events with an electrode placed in mesencephalon we observed no change in seizure event frequency during a 60 min 4-PBA (50 M) treatment (**Fig. 1H**). While 4-PBA was shown to alleviate neuronal dysfunction and locomotor deficit in Munc18-1 worms, this treatment required a much higher concentration (5 mM) with associated toxicity concerns. Testing non-toxic concentrations of 4-PBA in *stxbp1* zebrafish models, however, failed to alleviate motor deficit or seizure activity phenotypes.

To further expand and explore additional pharmacological treatments for movement disorders seen in STXBP1-related disorders, we used 4-PBA as a ligand for AI-based screening of a 2315 compound approved drug library. Ligand-based virtual screening yielded 13 candidates (**Table 1**). To this list we added 3 pharmacological chaperones (Compounds A, C and D) identified in a previous virtual screen of 255,780 compounds and tested in the Munc18-1 (*STXBP1*) *C. elegans* C. model (Abramov et al., 2021). Candidate drugs were administered for 24 hr (4-5 dpf) at concentrations of 25 or 50 µM with individual larvae arrayed in a 96-well plate; locomotion assays were performed at 5 dpf. Heartbeat presence and response to touch were used to monitor toxicity. Tetracaine data from WT (n = 132) and *stxbp1a*^-/-^ (n = 96) zebrafish were aggregated to calculate a mean ‘total distance traveled’ value of 279 mm (see **Fig. 1A**). As larvae do not move under tetracaine anesthesia, this value represents “false positive” movement data in single-point locomotion tracking and was used to normalize total distance traveled values for all drug candidates. In *stxbp1a* mutants, none of the drug candidates significantly altered movement (**Fig. 2 *stxbp1a***; **Supplemental Table 1**). Conversely, in WT larvae, three compounds significantly reduced total distance traveled compared to on-plate non-treated controls. These include an adrenergic receptor agonist, guanfacine at 50 µM (**Fig. 2B**) and two histamine receptor antagonists with sedative potential, chlorpheniramine at 25 µM (**Fig. 2E**) and pheniramine at 25 and 50 µM (**Fig. 2F**). The remaining drug candidates did not induce significant changes in total distance traveled in WT larvae (**Fig. 2 WT**; **Supplemental Table 1**). However, following a Bonferroni correction for multiple comparisons, no drug significantly altered locomotion in either genotype suggesting these nominally significant locomotion changes in WT larvae reflect ‘false positives’. Taken together, these studies extend our observations with 4-PBA and do not support a therapeutic effect on *stxbp1a* movement disorder comorbidity.

**Figure 2.**
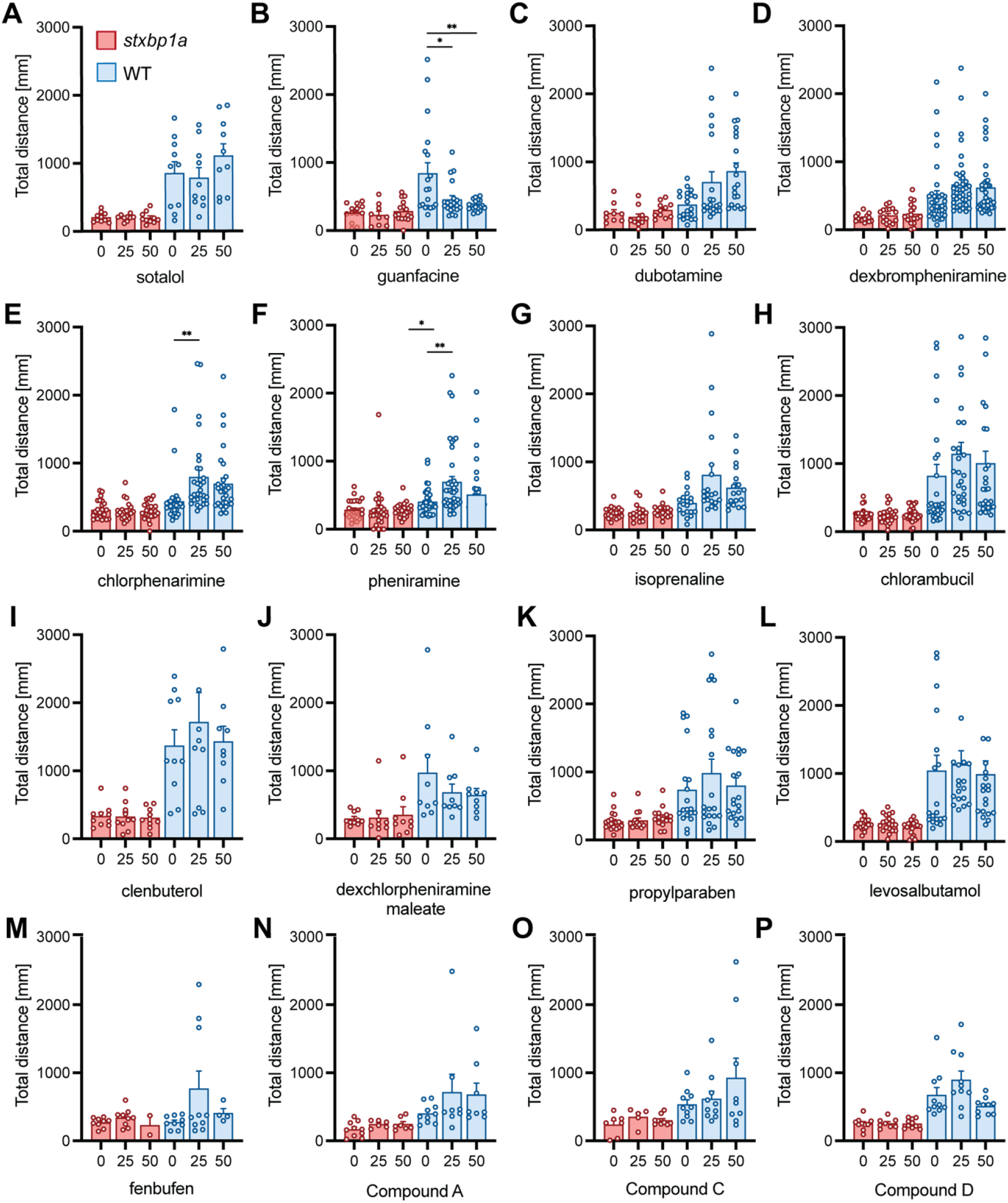
Screening of virtual drug candidates. (**A-P**) Bar plots of total distance traveled during a 10 min tracking epoch. Data is shown for *stxbp1a* mutant (salmon) and WT siblings (blue). Each dot represents one fish, comparing each condition to the non-treatment group (WT). N = 10-30 fish for each concentration and each drug. Date presented as mean S.E.M. Significant differences are denoted as **p≤0.01, *p≤0.05. One-way ANOVA with Dunnett’s multiple comparison test.

At the conclusion of locomotion assays, all larvae were again individually assessed under a dissecting microscope for signs of toxicity e.g., heartbeat presence and responsiveness to touch. Most larvae survived treatment drug candidates irrespective of genotype (**Fig. 3**). However, propylparaben at 25 µM (50% for *stxbp1a*, 25% for WT) and compound A at 25 µM (40% for *stxbp1a*, 10% for WT) induced considerable toxicity. These results suggest that unlike the potentially toxic mM drug concentrations used in Munc18-1 worms to demonstrate efficacy, drug concentrations used in our zebrafish assays were largely outside a range that can induce noticeable toxicities. Therefore, when considering these varied results from preclinical models, it is advisable to interpret with caution as safe concentrations of 4-PBA and similar compounds may not be adequate to achieve therapeutic effects.

**Figure 3.**
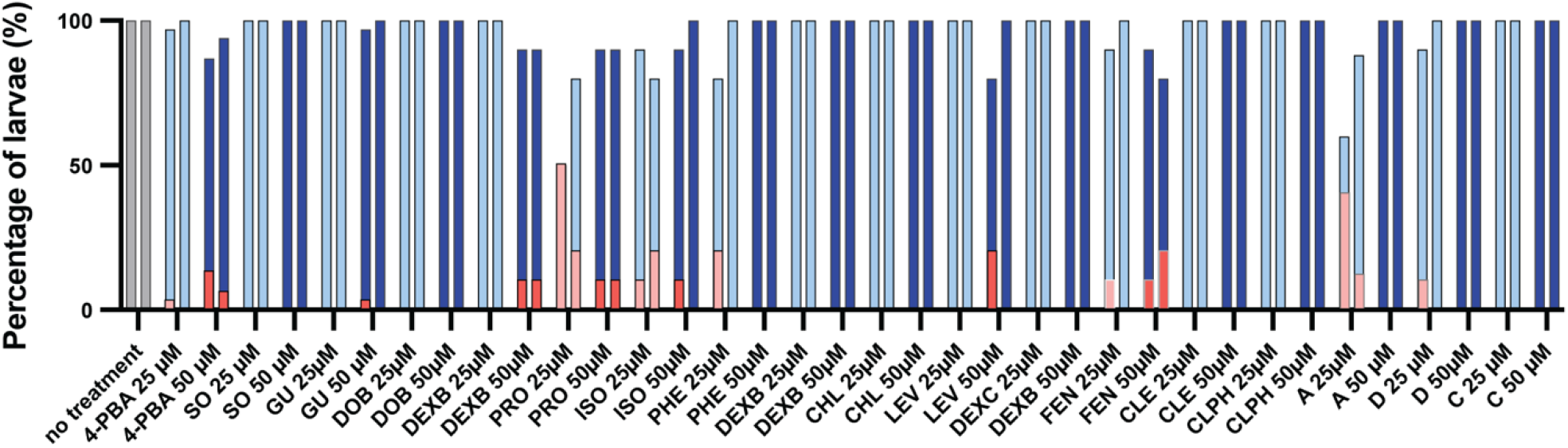
Toxicology assay for larvae following behavioral assessment. Toxicity assessment of all compounds at both concentrations. The left bar for each concentration represents *stxbp1a* mutants, while the right bar represents WT zebrafish. Light blue and red bars indicate survival and toxicity percentages at 25 µM, respectively, while dark blue and red bars indicate survival and toxicity percentages at 50 µM concentration, respectively. The “no treatment” condition serves as the baseline control with 0 concentration. Survival percentages for both WT and *stxbp1a* are provided, with overlaid percentages depicting larvae that did not survive treatment. N = 30-60 fish per condition. N (no treatment) = 300 WT and 270 *stxbp1a* larvae; N = 899 *stxbp1a* total, N = 990 WT total. Abbreviations: propylparaben (PRO), dexchlorpheniramine maleate (DEXC), isoprenaline (ISO), levosalbutamol (LEVO), clenbuterol (CLE), pheniramine (PHE), sotalol (SO), fenbufen (FEN), dobutamine (DOB), guanfacine (GU), chlorpheniramine (CLPH), dexbrompheniramine (DEXB) and chlorambucil (CLO).

## Discussion

Drug repurposing for rare genetic disorders is emerging as a promising strategy to identify new treatments for these medically intractable patients. Although more efficient than rational drug design-based strategies or simple high-throughput screening of existing novel chemical entities, repurposed drug development already incorporating known toxicity and tolerability profiles can still take up to 10 years and hundreds of millions of dollars. Zebrafish, with the capacity to generate hundreds of larvae per experiment, are a vertebrate species well suited for large-scale repurposed drug screening strategies. Artificial Intelligence-based virtual screening programs, which can be completed in minutes to hours, further expand our ability to accelerate this process (Wu et al., 2024, Gangwal and Lavecchia, 2024, Kim et al., 2020). In this study, we used AI-assisted screening of an approved drug library to identify drug candidates that we could test using sensitive phenotype-based assays in larval zebrafish.

4-phenylbutyrate is an FDA-approved drug for treatment of urea cycle disorders (Glinton et al., 2023) that has recently received considerable attention as a potential repurposed drug candidate for STXBP1-related disorders (Shen et al., 2024, Nwosu et al., 2022, Grinspan ZM, 2024). Using stxbp1 zebrafish models representing epilepsy (*stxbp1b*) and motor impairment (*stxbp1a*) phenotypes seen in these patients, we could uncover no evidence for therapeutic rescue of either phenotype with 4-PBA or any of 16 other drug candidates. Although Kang and colleagues reported that 4-PBA can mitigate spike-wave discharge seizures in *Slc6a1*^*+/S295L*^ and *Gabrg2*^*+/Q390X*^ mice (Shen et al., 2024, Nwosu et al., 2022) we failed to observe any changes in generalized ictal-like seizure discharge frequency in *stxbp1b* mutant zebrafish. Rescue of the body bend locomotion deficit seen in Munc18-1 worms was reported by Burre and colleagues (Guiberson et al., 2018) at a concentration (5 mM) which may be toxic and was not replicated here in *stxbp1a* mutant zebrafish featuring a severe spontaneous swim movement deficit. While it is not always possible to resolve discordant findings from different experimental animal models (or from different laboratories)(Steward, 2016, Landis et al., 2012, Prager et al., 2019), these discrepancies warrant further investigation in scientifically rigorous cross-species studies. Seizure frequency and gross deficits in motor development were recently proposed as primary endpoints in clinical trials for STXBP1-related disorders (Xian et al., 2023). However, our preclinical findings using nearly 1900 zebrafish faithfully replicating these two primary endpoints do not support pursuit of 4-PBA or (related pharmacological candidates) for treatment of seizures or motor deficit in these patients.

## Methods

### Zebrafish Husbandry

All experimental procedures adhered to guidelines set by the National Institutes of Health and the University of California, San Francisco, and approved by the Institutional Animal Care and Use Committee (IACUC approval; #AN171512-03A). Adult zebrafish and larvae were maintained under controlled temperature conditions with a 14-hr light and 10-hr dark cycle. Adults were fed twice daily with a diet consisting of tropical flakes (Tetramin) and live brine shrimp. Larvae and embryos were raised in polystyrene petri dishes (100 × 20 mm; FisherBrand #FB0875711Z) in an incubator at 28°C, under the same light/dark cycle as the facility, in an “embryo medium” comprising 0.03% instant ocean (Aquarium System, Inc.) in reverse osmosis-distilled water. Wild-type (WT) AB strain and stable germline mutations in *stxbp1* that had been backcrossed to AB WT for at least 10 generations were used for all studies. Homozygous *stxbp1a* mutant (n = 899) or WT larvae (n = 990) were used for behavioral experiments; homozygous *stxbp1b* mutant (n = 9) larvae were used for electrophysiology experiments. Larvae between 3- and 7-days post-fertilization (dpf) lack discernible sex chromosomes.

### Genotyping

Genomic DNA was extracted from whole larvae using Zebrafish Quick Genotyping DNA Preparation Kit (Bioland Scientific). We amplified *stxbp1a* gDNA using the following primers: stxbp1a-F: CACACACTTACAGCAGGAATGAGTGG, stxbp1a-R: ATTCAGAC

TTCAACTGTACATGTATTGTG. These primers amplify a 275-bp region including the *stxbp1a* mutation site. The mutant allele was then detected by digesting the amplicon with BsaHI, for which the restriction site is absent in *stxbp1a* mutants, and electrophoresis to separate the digested samples on a 2% agarose gel. We amplified *stxbp1b* gDNA using the following primers stxbp1b-F: ATC TGC GTA GAA AGC TGA GCT TCA TAG, stxbp1b-R: GTA ATGAAAGCCTATGCACCA. Mutant alleles were detected by digesting the amplicon with BsiHKAi, for which the restriction site is absent in *stxbp1b* mutants, and electrophoresis to separate digested samples on 2% agarose gel (Grone et al., 2016).

### Virtual Drug Screening

DrugRep is an automated, parameter free computer-aided drug discovery online tool for virtual screening of drug libraries (Gan et al. 2023). For *in silico* screening using DrugRep, we employed a ligand-based approach using the chemical structure for 4-phenylbutyrate (4-PBA). We screened an approved drug library (n = 2315 drugs) sourced from the DrugBank database (version 5.1.7) enhanced for clinically approved drugs; Flexi-LS-align was selected for its enhanced structural flexibility (Hu et al., 2018). This approach accommodates various conformers derived from 4-PBA, demonstrating consistent high similarity scores (>0.9) across multiple drug classes, minimal toxicity concerns, and a narrow LogP value range (Table 1). Compounds with LogP values outside the range of 0 to 5 and identified toxicity concerns, such as orciprenaline, terbutaline, and lofetamine l-123 were excluded.

### Locomotion Assay

At 3 dpf, *stxbp1a* mutant larvae were identified as unhatched embryos (Grone et al., 2016) and manually released from chorion using fine forceps. At 5 dpf, WT and *stxbp1a* mutant zebrafish larvae were placed in a 96-well plate with one larva per well and 5-10 larvae per treatment group. The larvae were placed in 150 μL of solution containing embryo media or test drug. Larvae were exposed to drug for 48 h in an incubator in the dark at 28.5 0.5 C. Solutions were changed every 24 hr. Following drug exposure, zebrafish larvae underwent a repeat of the toxicological evaluation described below. Plates were then acclimated in the observation chamber for 20 min. A DanioVision system, capturing video at 25 frames per second and running EthoVision XT 11.5 software (DanioVision, Noldus Information Technology), was used for automated single-point detection of larval movement. Video acquisition was performed with the following settings: subject: darker than background, method: differencing, sensitivity: ∼25-30, background changes: video pixel smoothing and fast subject contour: 1 pixel with contour dilation set to erode first then dilate, subject size: minimum 15 and maximum 5178. A baseline 20 min acquisition was performed (Trial 1). Control wells containing embryo media were then replaced with media supplemented with tetracaine (200 μM) to obtain a ground truth measurement of the DanioVision system tracking error e.g., larvae do not move when bathed in a local anesthetic. Following a 15 min incubation, a second 20 min ‘recording’ (Trial 2) acquisition was performed under the same parameters. The locomotion assay was performed in the dark to minimize risk of drug photodegradation. All assays were conducted between 9:30 AM and 12:00 PM P.S.T. All screening experiments were performed on at least two biological replicates.

### Toxicology Assay

Individual larvae were examined under a Leica dissecting microscope. Toxicity assessment included observation of a visible heartbeat, intact circulation, and response to gentle touch (Moog & Baraban, 2022; Whyte-Fagundes et al., 2024; Lee & Yang, 2021; von Hellfeld et al., 2023). For *stxbp1a* mutant larvae, characterized by immobility, presence of a visible heartbeat and/or intact circulation were evaluated. Solution was changed, and toxicity was assessed at 24- and 48-hours post-incubation, before initiating baseline locomotion assay (Trial 1) and following a test drug recording (Trial 2). Results from each toxicity assay were recorded, and deceased fish were excluded from subsequent analysis.

### Pharmacology

Compounds used in the study were commercially sourced from Millipore-Sigma (propylparaben, PRO; dexchlorpheniramine maleate, DEXC; isoprenaline, ISO; levosalbutamol, LEVO; clenbuterol, CLE; pheniramine, PHE) or VWR (tetracaine, TET; sotalol, SO; fenbufen, FEN; dobutamine, DOB; guanfacine, GU; chlorpheniramine, CLPH; dexbrompheniramine, DEXB; chlorambucil, CLO). Compounds A, C and D were sourced from Molport. Fenbufen, compounds D, C, and A were initially dissolved in 100% DMSO (dimethyl sulfoxide, Sigma Aldrich, CAS: 67-68-5) to a final stock concentration of 1 mM, then diluted in embryo medium to achieve assay concentrations of 25 μM (with 0.025% DMSO) and 50 μM (with 0.05% DMSO). All other compounds were dissolved directly in embryo media (E3) and further diluted as required. Final solutions were freshly prepared and used within seven days. Four compounds (neomycin A, ZINC9180966, ZINC32124456) were excluded from screening due to their insolubility in both embryo media and DMSO. All compounds are listed in Table 1 with their LogP solubilities.

### Analysis

Locomotion data were extracted from recorded videos, and toxicity was used to exclude empty wells and/or embryos. The locomotor activity for each 20 min recording session was then analyzed by calculating mean “total distance travelled (in mm)” extracted from EthoVision XT 11.5 (Wageningen, The Netherlands) imported into Microsoft Excel 16.9 (Seattle, WA) and visualized using GraphPad Prism 10.4 software (Boston, MA). A Z’-factor was calculated as follows: Z’-factor = 1–[3*(σp+σn)/◼μp–μn◼] where μ represents mean value of the baseline (WT with embryo media) and σ denotes standard deviations of baseline and negative control for movement (200 µM tetracaine). Pooled data was used to calculate a Z’-factor for our locomotion assay. Resulting in Z’-factor of -5.936 which is in the range (−7.1 to 0.78) considered satisfactory and consistent with published phenotypic larval zebrafish screens (Lescouzères et al., 2023; Liu et al., 2016).

To mitigate background noise inherent in the DanioVision single-point detection algorithm, zebrafish in the tetracaine control group were aggregated. The mean of these trials, 279 mm total movement, served as an internal control group and baseline for non-specific detection of immobile larvae. Trial locomotion data was then normalized by dividing raw score on total distance by 279 mm.

### Statistics

Statistical analyses were conducted using GraphPad Prism. Assessment of normality of the distribution of data was determined with the Shapiro-Wilk test, to apply either a parametric or nonparametric test. An unpaired t-test with Welch’s correction or One-way analysis of variance with Dunnett’s multiple comparison test was utilized to assess significance. Differences between experimental groups are deemed significant for *p ≤ 0.05; **p ≤ 0.01; ***p ≤ 0.001; ****p ≤ 0.0001. All individual values were reported alongside data distribution or as mean ± standard error of the mean (S.E.M.). To control for multiple comparisons across all drug candidates, Bonferroni correction was applied to all post-hoc p-values across the full family of 70 comparisons (corrected threshold: p < 0.000714). Every single *stxbp1a* post-hoc comparison has p ≥ 0.18, with the vast majority above 0.80. There is no borderline signal in the mutant genotype across any drug, concentration, or compound class tested.

## Abbreviations

(CRISPR): clustered regularly interspaced short palindromic repeats
(STXBP1): dpf (days post-fertilization);syntaxin-binding protein 1
(PTZ): pentylenetetrazole
(4-PBA): 4-phenylbutyrate

## Data availability

This study includes no data deposited in external repositories.

## Author contributions

A.F., P.W-F. and S.C.B. conceived and designed experiments; A.F. performed behavioral drug screening assays; P.W-F. performed electrophysiology assays; A.F., P.W-F. provided edits on draft versions of the manuscript; S.C.B. wrote the manuscript.

## Acknowledgements

This work was supported by grants from the STXBP1 Foundation and NIH R01-NS096976, R01-HD102071, R21-NS138525 and U54-NS117170 (to. S.C.B.) and a postdoctoral fellowship from the Savoy Foundation of Canada (to P.W-F). We thank Julia Schles and Eoghan Trainor for maintenance of zebrafish lines.

